# Multi-parametric assays capture sex- and environment-dependent modifiers of behavioral phenotypes in autism mouse models

**DOI:** 10.1101/2024.01.04.574201

**Authors:** Lucas Wahl, Arun Karim, Amy R. Hassett, Max van der Doe, Aleksandra Badura

## Abstract

Current phenotyping approaches for murine autism models often focus on one selected behavioral feature, making the translation onto a spectrum of autistic characteristics in humans challenging. Furthermore, sex and environmental factors are rarely considered.

Here, we aimed to capture the full spectrum of behavioral manifestations in three autism mouse models to develop a “behavioral fingerprint” that takes environmental and sex influences under consideration. To this end, we employed a wide range of classical standardized behavioral tests; and two multi-parametric behavioral assays: the Live Mouse Tracker and Motion Sequencing (MoSeq), on male and female *Shank2*, *Tsc1* and Purkinje cell specific*-Tsc1* mutant mice raised in standard or enriched environments. Our aim was to integrate our high dimensional data into one single platform to classify differences in all experimental groups along dimensions with maximum discriminative power. Multi-parametric behavioral assays enabled far more accurate classification of experimental groups compared to classical tests, and dimensionality reduction analysis demonstrated significant additional gains in classification accuracy, highlighting the presence of sex, environmental and genotype differences in our experimental groups. Together, our results provide a complete phenotypic description of all tested groups, suggesting multi-parametric assays can capture the entire spectrum of the heterogenous phenotype in autism mouse models.

## Introduction

Mouse models serve as invaluable tools in the comprehensive exploration of human neurodevelopmental disorders, and have been used to understand the underlying pathophysiology, evaluate new mechanistic targets, and screen novel drugs^1–5^. However, one of the limiting factors of using mouse models for complex neurodevelopmental disorders relates to the pronounced levels of phenotypic variability often observed in humans, which is difficult to study in the laboratory setting, that frequently resorts to tests that measure a limited number of behaviors^6^.

Autism is a prime example of such a complex neurodevelopmental disorder with a broad spectrum of behavioral manifestations^7^. This heterogeneity comes, in part, due to the etiology of autism which has a large genetic component^8,9^ but whose risk is also highly sensitive towards environmental modifiers^10^. In part due to this extensive phenotype variability, autism diagnosis in humans is a lengthy process that encompasses direct observation of the individuals, which can be extended with neuropsychological evaluation tests, and interviews with parents or caregivers^11,12^. Therefore, there have recently been considerable scientific efforts to identify distinct subgroups within the spectrum of autism^13–15^.

Pre-clinically, laboratory animal models have been used to establish autism-like phenotypes primarily through the development of mouse models with high-risk genetic mutations^2,16,17^. To establish autism-like phenotypes in the autism mouse models, several behavioral paradigms have been developed, which have been commonly used to assess primary behavioral domains of core autism deficits^18^. While these classical behavioral assays were mostly designed for single-purpose quantification of specific behavioral impairments (such as perseveration, repetitive behaviors or social deficits), recent computational advances have allowed exploration of behavioral domains in significantly greater detail, enabling classification of behaviors in home cages^19^, studying group dynamics and defensive behaviors^20,21^, and providing automated quantification of complex social behaviors^20,22,23^. Additionally, methods that rely on unsupervised machine learning have provided tools to quantify so-called behavioral “modules”, “motifs” or “syllables”, that have been defined as sub-second building blocks of more complex motor sequences^24,25^. Together, these tools can effectively and reliably capture different aspects of the phenotypic diversity and have recently been introduced for the quantification of neurodevelopmental deficits in mouse models^26,27^.

Research using animal models of autism has focused mainly on the genetic mechanisms underlying this disorder. Several studies investigated home-cage enrichment as an environmental modifier, which has been shown to ameliorate the autism-like phenotype in some genetic models^28–30^. However, sex is also a modifier of autism risk. While historically the prevalence of autism showed distinct disparity in males and females, with a ratio of 4:1^31,32^, newer studies point to a ratio closer to 3:1 or 2:1^33^. Autism in females often presents differently from the stereotype of autistic behavior^34,35^ and is therefore frequently mis- or underdiagnosed^36,37^. Furthermore, most studies of autism consistently enroll small numbers of females or exclude them altogether, creating a “leaky pipeline” that results in severe underrepresentation of females^38^. Similar sex-bias is present in fundamental autism research using mouse models, where males are overrepresented^39^. It is therefore increasingly urgent to establish an autism phenotype in female mice representing the most commonly used genetic autism models, while taking the environmental modifiers into consideration.

Here, we combined phenotyping using classical behavioral assays with recent advances in automated classification of individual and group behaviors to develop a new dimensionality reduction approach that captured sex- and environment-dependent variability of behavioral phenotypes in *Shank2^-/-^* and *Tsc1^+/-^*autism mouse models, as well as Purkinje cell specific, heterozygous *Tsc1* mutant mice (further referred to as *L7-Tsc1^flox/+^* mice). *SHANK* genes, which play an essential role in the formation and the maintenance of synapses, have an established association with autism as monogenic risk-factors^40^. Loss-of-function mutations in the *TSC1* gene result in tuberous sclerosis complex (TSC), a genetic disorder with high levels of comorbidity for autism^41^. Additionally, we used Purkinje cell specific mutants of *Tsc1*^42^, as cerebellar injury at birth has been described as one of the highest risk factors for autism^43^, indicating both the importance of normal cerebellar development and the impact of cerebellar-specific genetic deletion on autism-like phenotypes. Using classical behavioral assays, while *Shank2^-/-^* mice showed clear behavioral deficits compared to wildtype littermates, these were unable to capture genotype effects in *Tsc1^+/-^* and *L7-Tsc1^flox/+^* mice. Contrastingly, multi-parametric assays were able to capture sex- and environmental-dependent effects onbehavioral autism-like phenotypes in all models. Machine learning classification of combined outputs of classical and multi-parametric behavioral assays showed high levels of accuracy where single behavioral assays failed to detect differences in experimental groups.

Our approach can be applied to a range of behavioral data outputs and neurodevelopmental and neuropsychiatric animal models to show meaningful lower-dimensional behavioral dynamics that allow quantitative description of heterogeneous mouse behavior.

## Results

### Classical behavioral assays capture modulating effect of environmental enrichment on autism-like phenotypes in Shank2^-/-^ mice

In order to test environmental effects and sexual behavioral dimorphism on a wide range of autism-like phenotypes, male and female mice were assigned to either standard housing or housing containing environmental enrichment at weaning age (**fig. 1A**). At 6 weeks old, animals were injected with RFID tags to enable group tracking using the Live Mouse Tracker system (LMT^20^). Next, all animals underwent a series of classical behavioral assays as well as Motion Sequencing (MoSeq)^24^, to capture a broad range of behavioral phenotypes.

**Figure 1.**
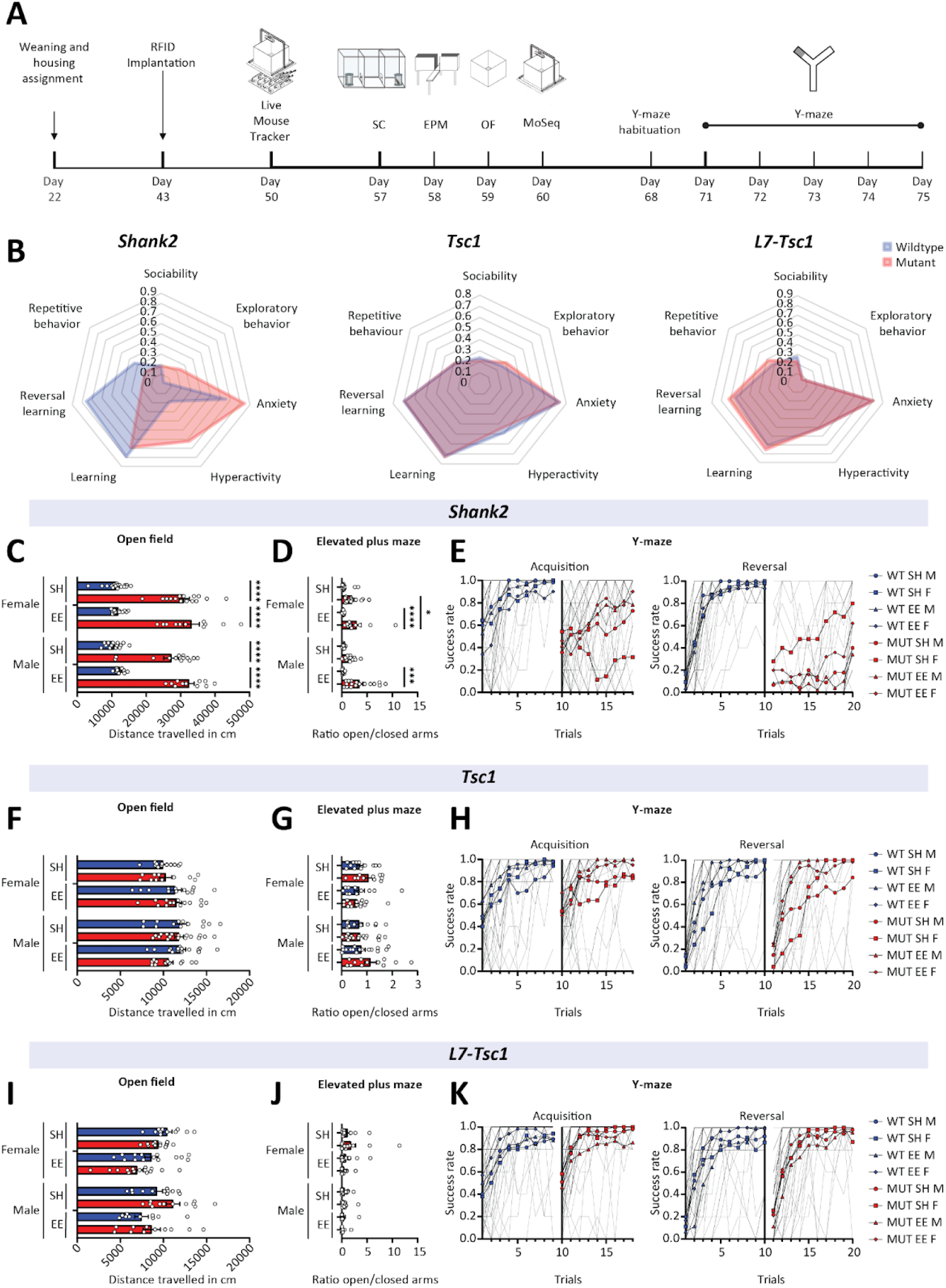
Behavioral phenotyping of autism mouse models using classical behavioral assays. **A)** Experimental timeline of behavioral experiments. SC, social chamber; EPM, elevated plus maze; OF, open field; MoSeq, motion sequencing. **B)** Normalized scores in classical behavioral assays per mouse model of autism. **C, F, I)** Distance traveled in the open field test. Data presented as mean with SEM (three-way ANOVA). **D, G, J)** Ratio of the time spent in the open arms of an elevated plus maze divided by the time spent in the closed arms. Data presented as mean with SEM (three-way ANOVA). **E, H, K)** Performance in the acquisition phase of the Y-maze (left) and the reversal phase (right). Data presented as mean with SEM (one-way ANOVA). * p ≤ 0.05, ** p ≤ 0.01, *** p ≤ 0.001. n = 325 mice.

In the classical behavioral assays, *Shank2^-/^*^-^ mice were found to show behavioral impairments in anxiety related behavior (**fig. 1B, supplementary fig. 1A**) as well as a pronounced hyperactivity phenotype (genotype: *p* < 0.0001; **fig. 1C**) in the open field test, and deficits in learning (*p* < 0.0001) and reversal learning (genotype: *p* < 0.0001) during the water y-maze when compared to their wildtype littermates (**fig. 1E**). *Shank2^-/^*^-^ mice showed increased levels of exploratory behavior (genotype: *p* < 0.0001; **fig. 1D**) in the elevated plus maze, an effect which was exacerbated in animals raised in enriched environments (genotype x housing: *p* = 0.0076). *Shank2^-/^*^-^ mice showed similar levels of social behavior compared to their wildtype littermates, and reduced burying behavior (genotype: *p* < 0.0033; **supplementary fig. 1B-C**). Neither the *Tsc1^+/-^* (**fig. 1B**, **center**) nor the heterozygous *L7-Tsc1^flox/+^* (**fig. 1B, right**) mice showed significant impairments in the classical behavioral assays (**fig. 1F-K, supplementary fig. 1**).

### Motion Sequencing reveals genotype, sex and environment-specific behavioral phenotypes in mouse models or autism

Next, we performed Motion Sequencing (MoSeq) during an extended open field test, to study the underlying behavioral state of the animals. MoSeq uses unsupervised machine learning to find repeated behavioral patterns referred to as behavioral syllables^24^. A key difference to most other behavioral assays is that usually behavioral testing is done in response to a specific factor, such as a novel mouse, novel object or a learning task. Here, we aimed to look at the innate behavioral differences expressed by the animals by studying the expression of behavioral patterns in an arena without external stimuli such as objects, tasks, or the presence of mazes, to see whether the autism-like phenotype can be segmented during freely expressed behavior.

A total of 40 behavioral syllables were identified through the MoSeq machine learning algorithm (**fig. 2A, C, E**). In *Shank2^-/^*^-^ mice, differential expression of behavioral syllables showed significant alterations in their usage depending on the genotype (**fig. 2B, top**) and sex (**fig. 2B, center**). However, no syllables were found to be differently expressed by *Shank2^-/^*^-^ mice raised in different environmental conditions (**fig. 2B, bottom**). This was surprising given that the classical elevated plus maze clearly showed significant differences in behavior dependent on environmental conditions. In the *Tsc1^+/-^* mice, MoSeq was able to detect sex specific expression of behavioral syllables (**fig. 2D, center**), yet no genotype- or environment-specific syllables were found (**fig. 2D, top and bottom**). Similarly, *L7-Tsc1^flox/+^*mice showed differential expression of sex specific syllables but no genotype specific syllables (**fig. 2F, top and bottom**). However, behavioral syllables specific to environmental conditions were clearly displayed in *L7-Tsc1^flox/+^*mutant mice (**fig. 2F, bottom**). These results indicate the ability of the MoSeq approach to accurately capture sex differences in all tested models, with some added predictive power for genotype differences in *Shank2^-/^*^-^ mice and environmental effects in heterozygous *L7-Tsc1^flox/+^* mice.

**Figure 2.**
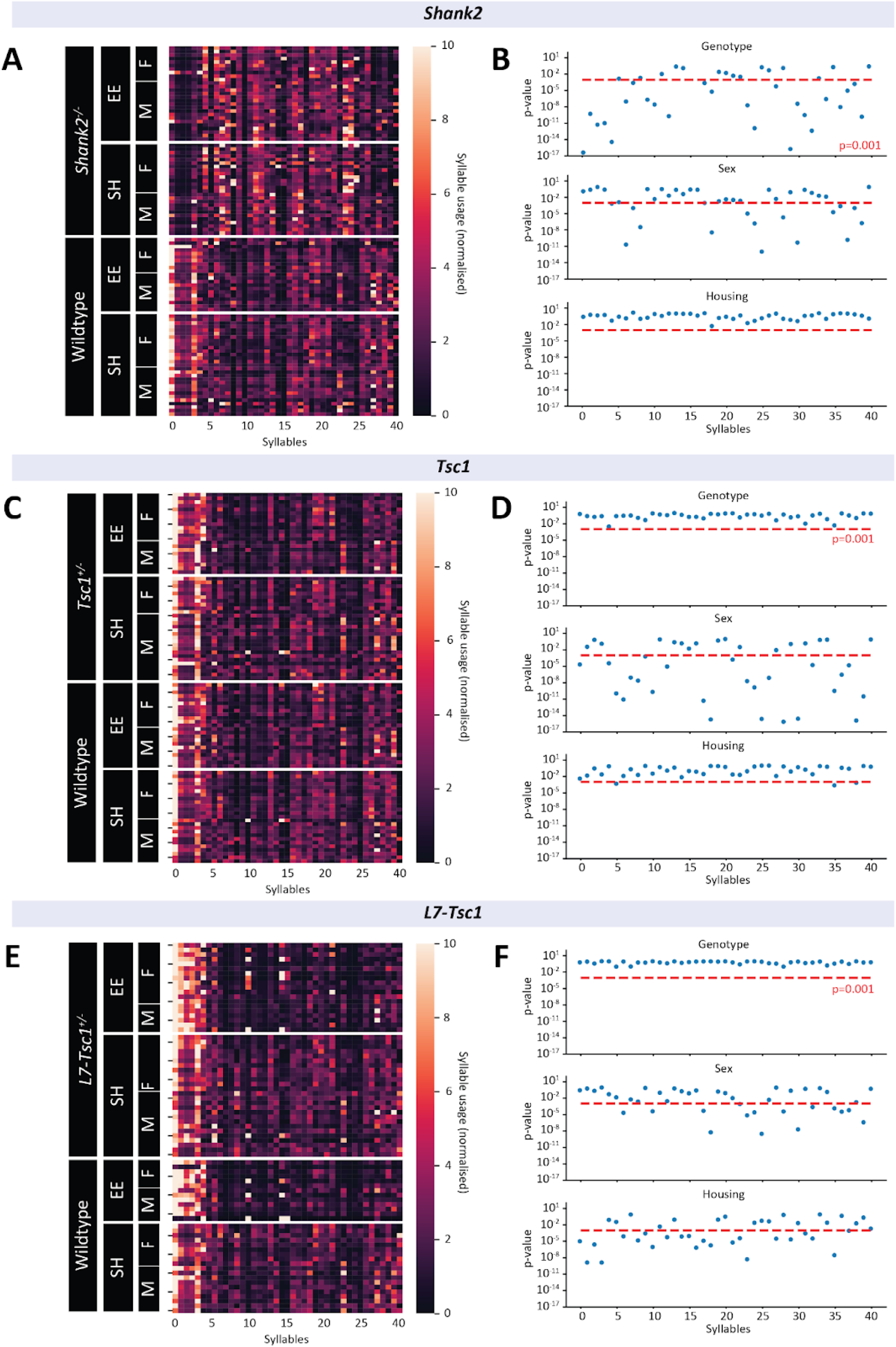
Unsupervised segmentation of behavior of autism models in an open field arena using motion sequencing. **A, C, E)** Heatmaps of normalized syllable usages per genotype. Each row represents an individual animal, each column represents a behavioral syllable. SH, standard housing; EE, environmental enriched housing; F, female; M, male. **B, D, F)** Significance of syllables for sex, housing and genotype differences, obtained by 3-way anova. A p value of 0.001 was calculated based on the number of variables and applied to correct for multiple comparisons. Each dot represents a syllable. n = 278 mice.

### Automated profiling of group behaviors in autism models reveals sex- and environment-dependent motor and social phenotypes

The LMT assay^20^ allows for the automated quantification of group behaviors in a home-cage-like environment containing enrichment such as bedding and nesting material. Furthermore, this paradigm is optimized for testing of animals in groups, whereas the previously mentioned behavioral assays only quantify the behavior of single animals, allowing for more naturalistic social expression of behaviors in a group of littermates that were housed together. Similarly to the performance in classical behavioral assays, *Shank2^-/-^* mice exhibited increased motor related behaviors and decreased immobility related behaviors in the LMT, reflecting the hyperactive phenotype we observed in the open field test (**fig. 3A**). Similar to results in the social chamber test, where *Shank2^-/^*^-^ mice exhibited normal levels of social behavior (**supplementary fig. 1B**), expression of most social behaviors in the LMT was found to be intact (**fig. 3A**). Interestingly however, behaviors related to group dynamics were strongly dysregulated, indicating normal overall social levels but differential expressions of social bouts. *Shank2^-/^*^-^ mice showed significant genotype-, sex- and environment-specific differences in several group and individual behaviors when compared to wildtype littermates (**fig. 3B, supplementary fig. 3**). *Tsc1^+/-^* mice displayed less pronounced phenotypic differences compared to *Shank2^-/-^* mice. Differential expression of social behavior was found between male and female animals raised in standard housing, with female *Tsc1^+/-^* mice generally expressing higher sociability and male *Tsc1^+/-^* mice displaying significant deficits in social behavior (**fig. 3C**). Furthermore, environmental conditions significantly modulated several motor related behaviors and social behaviors in *Tsc1^+/-^* mice, whereas sex only showed minor impact on measured behaviors, suggesting that differences in this model could be primarily environment and to a lesser degree sex dependent (**fig. 3D, supplementary fig. 3**). In *L7-Tsc1^flox/+^* mice, no significant difference in behaviors were found for both genotype and sex (**fig. 3E-F, supplementary fig. 3**). However, animals raised in enriched environments showed a significantly different behavioral profile compared to standard housed *L7-Tsc1^flox/+^* mutant mice (**fig. 3E-F**).

**Figure 3.**
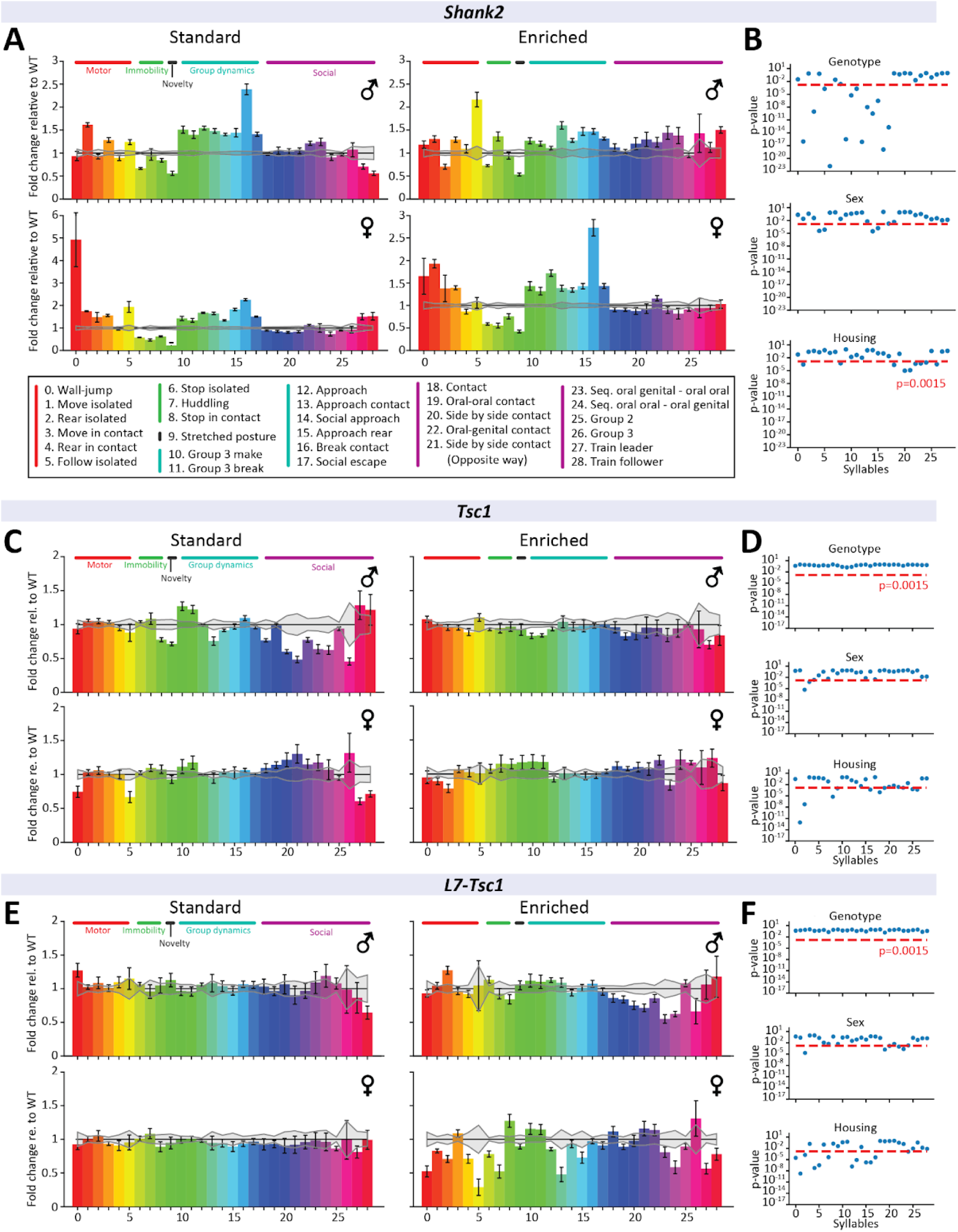
Behavioral analysis of autism mouse models in a home-cage environment. **A,C,E)** Fold change relative to respective wildtype results. Gray error lines indicate wildtype standard error. Data presented as mean with SEM. Each bar represents a behavior. **B, D, F)** Significance of behaviors for sex, housing and genotype differences, obtained by 3-way anova. Each dot represents a syllable. A p value of 0.0015 was calculated based on the number of variables and applied to correct for multiple comparisons. n = 266 mice.

### Combining classical and multi-parametric measurements achieves the highest classification accuracy across all mouse models

Classical behavioral assays, LMT and MoSeq have each shown the ability to quantify distinct aspects of autism-like behavioral phenotypes. To further examine the capabilities of classical and multi-parametric assays to capture the modulating effect of environment and sex on autism-like behavior, we performed PCA-LDA and machine learning classification on data collected across all assays. In total, 20 experimental measures were obtained from classical behavioral testing, information on 44 behavioral syllables was obtained from the MoSeq and 29 behavioral variables were obtained from the LMT. PCA-LDA reduced the dimensionality of the data from the initial 89 dimensions to N-1 dimension, where N is the amount of classes, resulting in 7 linear discriminants. 2-dimensional representation of the first 2 linear discriminants with the highest eigenvalue showed a clear separation between *Shank2* experimental groups (**fig. 4A**). The first linear discriminant best separates *Shank2^-/-^* mice and their wildtype littermates (**fig 4A**), explaining a substantial amount of the variability in the data (**fig. 4B**). Quantification of the variance explained per experiment shows that the majority of the explained variance between classes originates from MoSeq and LMT analyses (**fig. 4C**), indicating that these multi-parametric tests are highly capable of differentiating between *Shank2* experimental groups. To fully identify the ability of the classical and multi-parametric behavioral assays to differentiate sex, environmental conditions and genotypes, a linear support vector machine (SVM) classifier was trained for a multitude of combinations of classical and multiparametric assays. Individually, classical and multi-parametric assays only reached classification rates between 65-75%, while combining behavioral assays yielded substantial gains in classification accuracy (**fig. 4D**). Combining the various assays showed clear ability to accurately classify experimental groups in *Shank2^-/-^* mice and their wildtype littermates (**fig. 4E**). In *Tsc1^+/-^* and WT mice, visual representation of the first 2 linear discriminants shows clear segmentation between sexes by the first linear discriminant (**fig. 4F**), with a higher contribution to the total variability in the data of additional linear discriminants compared to *Shank2* experimental groups (**fig. 4G**). LMT analysis and MoSeq captured large amounts of the variability in experimental groups (**fig. 4H**). Linear SVM classifiers showed clear gains in classification accuracy by combining classical and multi-parametric behavioral assays (**fig. 4I**), in line with the more evenly distributed contribution of additional linear discriminants. This resulted in an accurate classification of most *Tsc1^+/-^* and WT experimental groups (**fig. 4J**). In *L7-Tsc1^flox/+^* mice and their wildtype littermates, the first 2 linear discriminants show strong separation between sexes and the different environmental conditions (**fig. 4K-L**). Similarly to the previous models, most of the variance in the experimental groups, which in this case is mostly notably related to variability due to environmental conditions, can be attributed to MoSeq and LMT analyses (**fig. 4M**). A combination of classical and multi-parametric assays showed clear increases in classification accuracy (**fig. 4N**), accurately classifying all of the experimental groups (**fig. 4O**).

**Figure 4.**
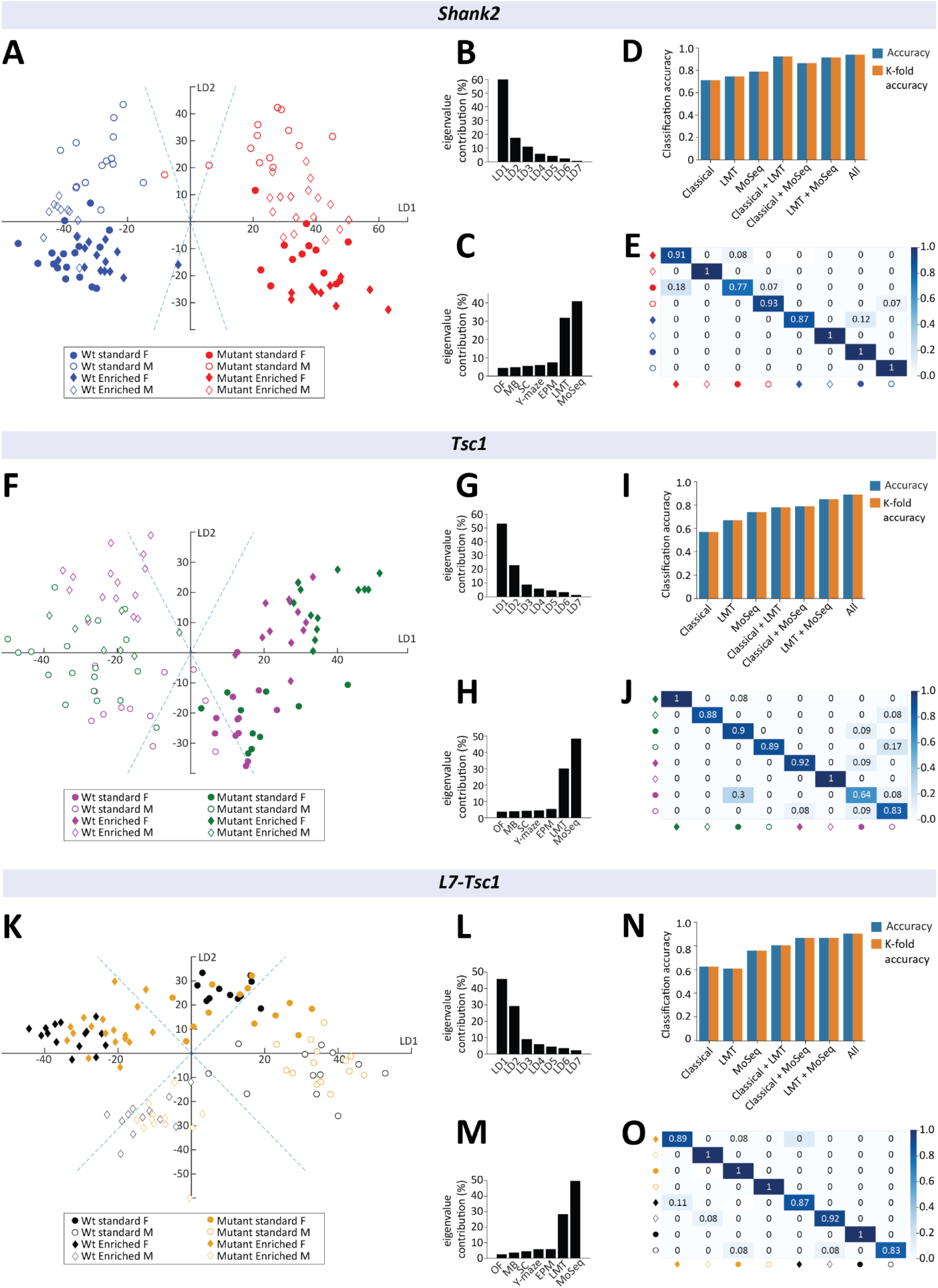
Classification accuracy of classical and multi-parametric data with reduced dimensionality. **A, F, K)** 2-Dimensional representation of the first 2 linear discriminants with the highest eigenvalue contribution, indicating the amount of variance captured. The x-axis represents the linear discriminant with the highest eigenvalue contribution (LD1), while the y-axis represents the linear discriminant with the second highest eigenvalue contribution (LD2). Standard housing conditions are identified by circles, while enriched housing conditions are represented using diamonds. Female and male mice are distinguished by filled and open symbols, respectively. **B, G, L)** Eigenvalue contribution of all linear discriminants, indicating the captured variance. **C, H, M)** Eigenvalue contribution of separate behavioral assays. SC, social chamber; EPM, elevated plus maze; OF, open field; LMT, live mouse tracker; MoSeq, motion sequencing. **D, I, N)** Accuracy and cross-validation for different combinations of behavioral assays used to train the linear SVM classifier. The y-axis shows the classification accuracy of the linear classifier and the accuracy result from leave-one-out cross validation. **E, J, O)** Confusion matrix of experimental groups made by incorporating behavioral data from classical and multi-parametric behavioral assays. The y-axis represents the true mice group the animals belong to, while the x-axis is the group which the classification model classified the mice as. Values in the confusion matrix represent the ratios of group prediction for the true groups. n = 328 mice.

Additionally, when behavioral data from all genotypes was combined (**fig. 5A-B**), classification accuracy showed a linear increase in performance with combined data from classical and multi-parametric assays (**fig. 5C**). Per-group classification shows especially high predictive validity between the genotypes, with higher confusion rates mostly restricted to comparisons within genotypes (**fig. 5D**). Together, these results indicate strong gains in classification accuracy of experimental groups by dimensionality reduction and a combination of both classical and multi-parametric behavioral assays.

**Figure 5.**
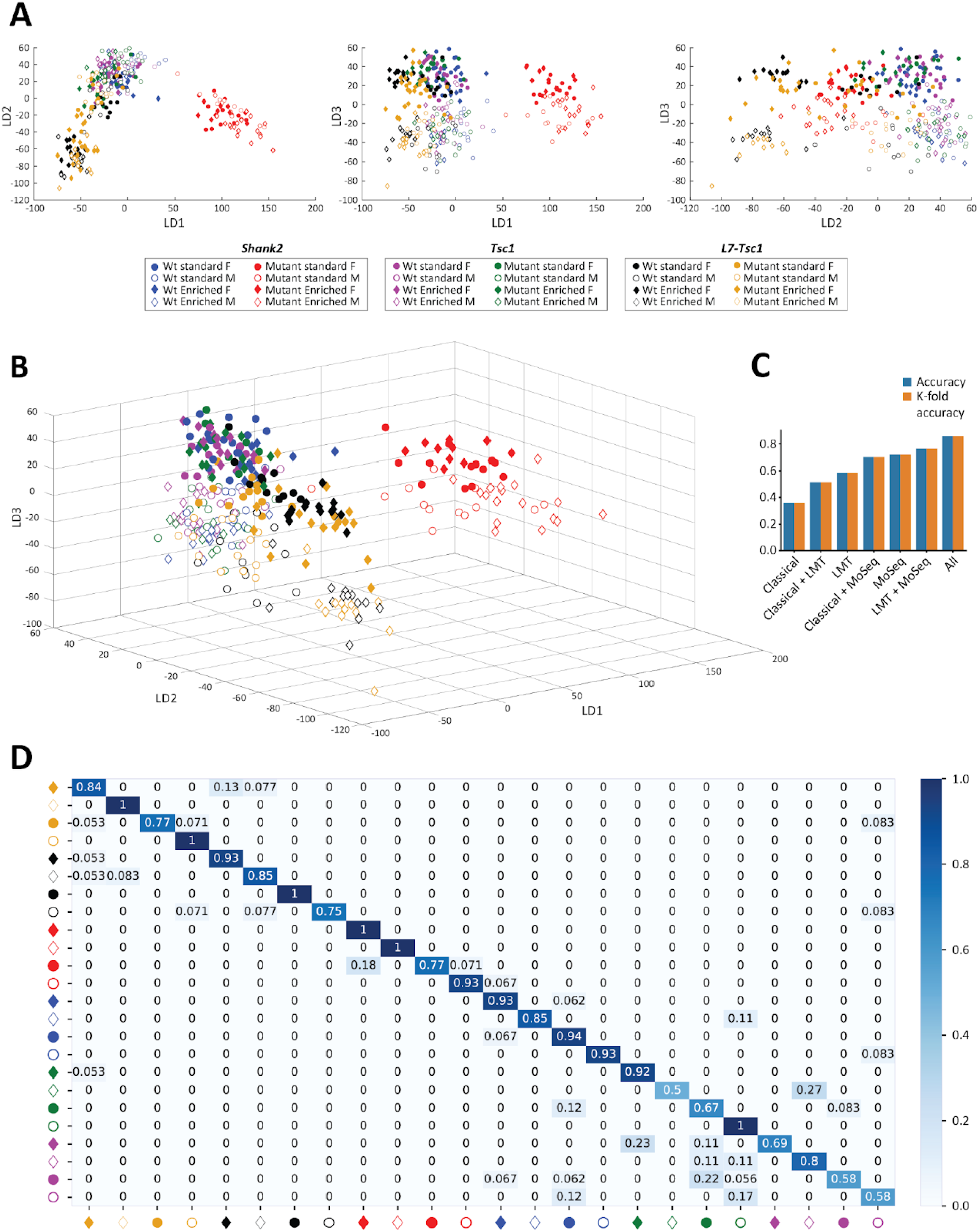
Classification accuracy for combined data of all genotypes. **A)** 2-Dimensional representation of the first 3 linear discriminants with the highest eigenvalue contribution, indicating the amount of variance captured. **B)** 3-Dimensional representation of the first 3 linear discriminants with the highest eigenvalue contribution. The x-axis represents the linear discriminant with the highest eigenvalue contribution (LD1) while the y-axis represents the linear discriminant with the second highest eigenvalue contribution (LD2) and the z-axis representing the linear discriminant with the third highest eigenvalue contribution (LD3). **C)** Accuracy and cross-validation for different combinations of behavioral assays used to train the linear SVM classifier. The y-axis shows the classification accuracy of the linear classifier and the accuracy result from leave-one-out cross validation. **D)** Confusion matrix showing all of the tested autism models and subgroups, made by incorporating behavioral data from classical and multi-parametric behavioral assays. The y-axis represents the true mice group the animals belong to, while the x-axis is the group which the classification model classified the mice as. Values in the confusion matrix represent the ratios of group prediction for the true groups. n = 328 mice.

## Discussion

Autism is a complex neurodevelopmental disorder which exhibits high genetic and phenotypic variability^7,12^, and it is therefore crucial to carefully study its diverse manifestations to understand the underlying biological mechanisms. Here we aimed to employ classical and multi-parametric assays to capture sex- and environment-dependent autism-like behaviors and explore the whole spectrum of phenotypic variability in three commonly used autism mouse models. Our study incorporates recent advances in automated behavioral analysis to demonstrate the power of integrating classical behavioral assays, MoSeq, and LMT, to comprehensively assess autism-like phenotypes in mouse models.

We observed that classical behavioral assays were very capable of accurately capturing several key aspects of previously reported behavioral deficits in *Shank2^-/-^* mice^44–46^. In our battery, *Shank2*-deficient mice exhibited anxiety-related behaviors, hyperactivity, and learning deficits (**fig. 1, supplementary fig. 1**). Moreover, growing up in enriched housing conditions significantly modulated the exploratory behavior of *Shank2^-/-^* mice in the elevated plus maze, emphasizing the significant impact of environmental conditions on behavioral phenotypes. Classical behavioral assays employed in the current study showed a lack of clear impairments in the *Tsc1^+/-^* and heterozygous *L7-Tsc1^flox/+^* mice which is in contrast with previous findings that showed social impairments in these lines^42,47^. We attribute this discrepancy to three possible factors: first, in contrast to the previous studies, all of our mouse lines were fully backcrossed (over 10 generations of crossing) into the *C57BL/6J* background whereas previous literature used *L7-Tsc1^flox/+^*mice on mixed background^42^. Because different background strains are known to have considerable phenotypic variability^48^, this could explain some of the observed differences. Second, since environmental factors have repeatedly been shown to affect behavioral outcomes ^49^, inherent differences in experimental conditions such as prior exposure to other behavioral tests, or differences in animal housing, temperature, light and experimenter handling may have contributed to the poor reproducibility of sociability deficits in *Tsc1^+/-^*and *L7-Tsc1^flox/+^* mice. Third, it is possible that for mice with relatively mild and variable phenotype like the *Tsc1^+/-^* and *L7Cre::Tsc1^flox/+^* failure to detect a significant difference between the genotypes in the classical assays can simply reflect the genetic heterogeneity within autism itself. Between 30% and 60% of people who harbor loss of function mutations in the *TSC1* and *TSC2* genes receive an autism diagnosis^50^ with mutations in the *TSC2* gene showing a higher risk^51^. It is therefore plausible that in those cases we need more refined tests to capture the diversity of behavioral phenotypes.

Recent trends in neuroscience have moved towards examination of more naturalistic behaviors^52^. By integrating laboratory experiments with less experimental constraints, more complex “natural” behaviors can now be quantified. There is an ongoing effort to develop analysis methods that properly quantify the behaviors underlying complex naturalistic movements^53^. Here we chose to employ two recently developed techniques - LMT^20^ and MoSeq^24^ - to assess their ability to discern subtle autism-like phenotypes, taking modifiers into consideration. Importantly both of those tools measure different properties: while the LMT enables identification of predetermined single and group behaviors for multiple mice using supervised learning, MoSeq dynamically builds representations of movement for individual mice utilizing unsupervised machine learning.

The automated profiling of home-cage-like behaviors using the LMT unveiled sex- and environment-dependent motor and social phenotypes. *Shank2^-/-^*mice exhibited a hyperactive phenotype consistent with classical assays and previous studies^44^. *Shank2^-/-^*mice also exhibited differential expressions of behaviors related to social group dynamics consistent with previous reports^44,45^, emphasizing the power of assessing social behaviors in a more naturalistic setting. Similarly, *Tsc1^+/-^* mice showed nuanced sex-dependent social behaviors and environment-dependent motor behaviors. *L7-Tsc1^flox/+^* mice exhibited distinct behavioral profiles under enriched housing conditions, further highlighting the interplay between genetic factors and environmental influences in shaping behavioral outcomes.

MoSeq, which in the past performed successful classification of pharmacological interventions^54^ and described deficits in a cerebellar-specific autism mouse model^26^, in our study was able to recognize genotype, sex, and environment-specific behavioral syllables in the tested mouse models. The identification of 40 distinct behavioral syllables revealed genotype- and sex-specific behavioral syllables in *Shank2*^-/-^ mice. However, no syllables were significantly affected by environmental conditions in these mice. It is possible that large genotype and sex differences in this particular mouse model are so dominant that they simply occlude other characteristics. *Tsc1^+/-^*mice exhibited sex-specific syllables, while *L7-Tsc1^flox/+^* mice displayed both sex and environment-specific syllables, highlighting the utility of motion sequencing in capturing nuanced aspects of behavioral variability specific to each model. The presence of sex-specific expression of behavioral syllables in all of the models we tested further reinforces the importance of considering sex differences in autism research^32^.

The integration of classical behavioral assays, MoSeq, and LMT allowed for a comprehensive assessment of autism-like phenotypes in all of the tested models. Machine learning classification further demonstrated the high predictive accuracy of a multi-parametric approach, in quantifying the behavioral impact of genotype, sex, and environmental conditions on autism-like phenotypes, emphasizing the synergistic value of integrating diverse behavioral assays for the classification of available mouse models of autism. Notably, we recognize that the preferred combination of assays may vary depending on the severity of the autism-like mouse models and the existence of environmental modifiers. For the quantification of more severe phenotypes, it could be sufficient to use specific, single-trait assays. However, for milder, more complex phenotypes, the combination of LMT and MoSeq might offer an in-depth description of the entire spectrum of the behavioral repertoire. This would allow higher throughput data collection than extensive single-trait behavioral assay batteries while offering a possibility to test interventions that aim at normalizing autism-like phenotypes.

In conclusion, our multi-parametric approach provides a nuanced perspective on the behavioral phenotyping of autism mouse models, offering insights into the interplay between genetics, sex, and environment. These findings contribute to the evolving understanding of the heterogeneity of autism and underscore the importance of comprehensive behavioral assessments in preclinical research. The integration of advanced methodologies opens avenues for more targeted investigations into the mechanisms underlying the diverse manifestations of autism-like phenotypes.

## Materials & Methods

### Experimental procedures

All experimental animal procedures were approved *a priori* by an independent animal ethical committee (DEC-Consult, Soest, The Netherlands), as required by Dutch law and conform to the relevant institutional regulations of the Erasmus Medical Center and Dutch legislation on animal experimentation (CCD approval: AVD1010020197846).

### Animals

A total of 325 mice were used in the described experiments (**supplementary fig. 2**). *Shank2^-/–^* (*^44^*; n = 59), *Tsc1^tm1Djk^* (*^55^*; referred to as *Tsc1^+/-^* in this manuscript; n = 54); and *CrB6.Cg-Tg(Pcp2-cre)3555Jdhu/J::Tsc1^flox/+^* (*^42^*; referred to as *L7-^-^Tsc1^flox/+^*in this manuscript; n = 58) mice models were used together with their wildtype littermates (n = 60, n = 54, n = 51, respectively). All mice were fully backrossed for >10 generations into the *C57BL/6J* background. Mice were weaned at P22, and group-housed (3 mice per cage) with mixed groups of wild-type and mutant littermates of the same genetic model. Male and female mice were housed separately. Mice were raised in either standard housing cages (SH; 335 cm^2^ floor space) with wood chip bedding (Lignocel) and nesting material or in environmentally enriched cages (EE; floor space of 800 cm^2^), which contained the same wood chip bedding and nesting material. Additionally, EE cages were provided with environmental enrichment in the form of a running wheel, wooden blocks, plastic tubes and paper houses. All mice were provided with food and water *ad libitum* and kept on a regular 12 h light/dark cycle.

### Order of experiments

For all mice, the first week of the behavioral battery consisted of an LMT recording. For the second week, the following schedule was kept at all times: Day 1: the social chamber test; Day 2: elevated plus maze; Day 3: open field test; Day 4: MoSeq. In the last week of the behavioral battery the water Y-maze was performed.

### Single-trait behavioral assays

All experiments except for the water Y-maze were performed in a behavioral box as described in^56^. This consisted of a 130 x 80 x 80 cm wooden box, which was lined with 6 mm high-pressure laminate and foam. The box contained white and infrared lights placed above a 10 mm Perspex® shelf. All video recordings of single-trait behavioral assays in the box were recorded with a fixed camera (acA 1300-600 gm, Basler AG) positioned above the arena. This camera was operated using the open-source software Bonsai^57^ with a frame rate of 25 frames per second (fps). Positional and locomotor analyses were done using the open-source software OptiMouse^58^. Animals were habituated to the testing room for 1 hour before experiments. All experimental arenas were cleaned with 70% ethanol between testing of different animals.

#### 1. Elevated plus maze

An elevated plus maze was utilized to examine anxiety-like behavior in mice^59^. The maze consisted of two open and closed arms, each with 29.5 × 8.5 cm white-opaque acrylic base, which were raised off the ground by cylindrical poles 30 cm in height. The closed arms were surrounded by black 20 cm high walls. Mice were placed in the center of a maze and left to explore for 10 minutes, after which they were returned to their home cage. The time spent in the open and closed arms of the maze, together with the locomotor activity and the number of transitions from the center of the maze to either closed or open arms, were quantified.

#### 2. Open field test

An open field test was used to examine anxiety-like behavior and locomotor activity ^60^. Animals could freely explore a 50 x 50 x 35 cm white-opaque acrylic arena for 30 minutes, after which the mouse was returned to the cage. The time spent in the inner and outer regions of the arena, together with the locomotor activity, were quantified.

#### 3. Social chamber test

A three-chamber social preference test was used to quantify sociability^61^. The apparatus consisted of a 63 × 42.5 × 21 cm transparent acrylic arena divided into three separate chambers. The divider walls contained an opening for the animals to pass through, which could be closed off with falling doors. First, the test animal was placed in the central chamber and allowed to freely roam for 10 mins for habituation to the apparatus. After this period, the doors were opened and the animal could freely explore all chambers for 10 minutes. Lastly, the experimental mouse was guided back into the middle chamber and two novel wire cups were placed in opposite chambers, with one containing a novel mouse. The experimental animal was then left to freely explore all chambers and the time spent near the novel mouse and empty cup was quantified, together with the total distance moved and number of transitions between chambers.

#### 4. Marble burying test

The marble burying test was performed as described previously^62^. In short, animals were transferred to a large cage (26,6 cm x 42,5 cm x 18,5 cm; Eurostand 1291H-type III H) filled with ∼4 cm of bedding and 20 evenly spaced out glass marbles and left for 30 minutes, after which the animals were removed. Burying behavior during the test was quantified using JAABA^62^.

### 5. Y-maze test

A five-day water Y-maze paradigm was used to examine cognitive flexibility^27,63^. A Y-shaped opaque polycarbonate apparatus consisting of symmetrical arms, each measuring 32 cm × 9 cm × 20 cm (length × width × height) from the center was filled with white, non-toxic, water-based paint (Basic color, 21 white 30081, Creall) to reduce visibility of an acrylic platform which would be placed in either arm. Notches in all three arms (8 cm from the center) allowed for placing a removable gate. Between each mouse, excrement was removed and water was exchanged to maintain an ideal water temperature between 22 and 26°C. To prevent distraction, a three-walled black shield was placed around the maze constructed of non-reflective black cardboard. The fourth wall was covered with a non-reflective black curtain.

Before experiments (Day 0), animals underwent habituation to the y-maze. During 3 habituation trials, animals were placed in each arm of the y-maze sequentially and were left to explore the maze for 60 s, after which the animal was removed. During the acquisition days (Days 1-2), a platform was placed at the end of either the left or right arm for 4 sessions of 5 consecutive 60 s trials. This location was kept the same for each mouse on the test day (Day 3), which consisted of a single session of 5 trials. Mice were required to have 80% success rate on the test day in order to progress to the reversal day, with the location of the platform switched to the arm opposite to the one presented in days 1-3. The reversal condition was introduced in the subsequent 2 days (Days 4-5). During the reversal day, mice were exposed to 4 sessions of 5 consecutive 60 s trials followed by a 5th “forced” session where a door was placed in the initial learned arm of the maze which no longer has a platform. All mice were kept in a clean cage under the heating lamp to dry in between the sessions and before returning to their homecage. Y-maze videos were recorded with a PsEye camera (Sony) at a frame rate of 30 fps. Correct choices, acquisition learning speed and reversal learning speed were measured. At the end of the day, water was removed, the maze was cleaned with 70% ethanol, and was left overnight to dry.

### Multi-parametric assays

Multi-parametric assays were performed to capture a wide range of behavioral profiles in an open field arena. We used the following two methods: the Motion Sequencing (MoSeq^24^) and the Live Mouse Tracker (LMT^20^). Both methods utilize unsupervised machine learning algorithms to automatically track mice and quantify multivariate behavioral profiles.

#### 1. Motion Sequencing (MoSeq)

MoSeq (v.1.1.1b0) was used as described previously^24,54^ to classify behavioral syllables, repeated sub-second movements patterns, through the use of an unsupervised machine learning algorithm. MoSeq was performed in single mice in a 50 x 50 cm opaque acrylic open-field cage placed inside a 120 cm wide, 120 cm deep and 160 cm tall behavioral box for a duration of 60 minutes. Mice were tracked by a Kinect2 (Microsoft) depth-sensing camera. Recordings were done at 30 fps using custom MoSeq acquisition software, described in Wiltschko et al. (2015). This data was then compressed and transferred to an analysis computer. The resulting data were processed using MoSeq analysis scripts, described in Wiltschko et al. (2015) and our custom written analysis pipeline [https://github.com/BaduraLab/ASD-mouse-model-analysis/tree/main]. In brief, the data files were uncompressed and pre-processed to identify the mouse in the arena. The mouse’s center of mass and orientation were detected, and the mouse was centered in an 80 x 80 pixel box. A custom-trained classifier was then used to rotate this box such that the nose of the mouse pointed rightward. This data was then used for subsequent processing steps. The MoSeq algorithm then performed principal component analysis (PCA) on the 80 x 80 pixel mouse data to reduce the dimensionality of the data. The mouse frame data was then projected onto the first ten principal components to yield a 10-dimensional time series that described the 3D pose dynamics of the mouse. This data was then used to train an autoregressive hidden Markov Model (AR-HMM) to cluster the mouse pose dynamic data into discrete behavioral modules (syllables). Finally, behavioral syllable transitions were computed by calculating the number of times each module was followed by each other module. We used 40 syllables with the highest usage over all mice to find the characteristic behavioral syllables of studied mouse models. Additionally, the total distance moved and the mean distance from the center were measured.

#### 2. Live Mouse Tracker (LMT)

The LMT was used to track mice group behavior. One week before experiments, mice were surgically injected subcutaneously with RFID tags (134 kHz Glass probe ISO 11784/11785m; Priority1Design) into the flank region, under isoflurane anaesthesia (induction: 4%V/V in O2, maintenance: 2-2.5% V/V in O^2^; flow rate ∼0.5 l/min) and subcutaneous injection of Rimadyl (5 mg/kg). The LMT set-up was built as described in de Chaumont et al. (2019) and placed into the same behavioral box used for MoSeq experiments. In short, the LMT consisted of a 50 x 50cm clear-walled enclosure, sitting atop a grid of 16 RFID coils (RFIDCOIL-100A.; Priority1Design). The Kinect camera was positioned above the enclosure and was controlled using the Live Mouse Tracker plugin of Icy^20^. Mice were recorded for a period of 60 minutes. LMT data consisted of a series of MP4 videos, for visual inspection of the experiment, and an SQLite database file containing behavioral data. The SQLite file was then pre-processed using a combination of custom-written Python and Matlab scripts and scripts made available by de Chaumont and colleagues. The LMT automatically labeled both individual and group behaviors, quantifying their frequency in real time using a deep learning algorithm. A set of 29 pre-determined behavioral events were quantified.

### Normalization

Data from single-trait behavioral assays was normalized to create radial plots showing an overview of the behavioral phenotype per autism mouse model. Measures used were: sociability, ratio of time spent near novel object divided by the time spent near novel object in the social chamber test; exploratory behavior, ratio of time spent in open arms divided by time spent in closed arms of the elevated plus maze; anxiety, time spent in inner zone of an open field test; hyperactivity, distance moved in an open field test; learning, area under the curve of the percentage of correct trials during the acquisition phase of the Y-maze; reversal learning, area under the curve of the percentage of correct trials during the reversal phase of the Y-maze; repetitive behavior, total time spent burying during the marble burying test.

To normalize the LMT data for each mutant group, we divided the time spent in each LMT behavior for the mutant mice with the mean time spent in each LMT behavior for WT mice with the same sex and housing conditions. This created a value of the fold change in behavior for the mutant mice relative to the WT mice.

### Dimensionality reduction & machine learning classification

The measurements from the single-trait behavioral assays and the multi-parametric assays were gathered in a singular dataset. Dimensionality of the data was reduced using a two-stage PCA-LDA algorithm, as Howland and Park (2004) have shown that using Principal Component Analysis (PCA) as a pre-processing step for Linear Discriminant Analysis (LDA) when performing dimensionality reduction allows for effective discrimination while successfully decreasing the dimensionality^64^. Variance captured by each individual single-trait and multi-parametric behavioral assay was calculated. To identify the capability of these assays to differentiate the different mice models and sex- and environmental-modifiers, the SciKitLearn (v0.24.2) python library^65^ was used to train linear classifiers. Because the hold-out approach, where the data is initially split between a training and test datasets for validation of the machine learning algorithm, creates larger uncertainties at smaller sample sizes^66^, we selected the Leave One Out (LOO) cross-validation to ensure that the classifiers were properly fitted. In short, LOO is a method for which the final model is trained on the entire dataset, allowing for higher accuracy in smaller datasets^66^, while still allowing for proper cross-validation through repeated linear classifier training within the dataset. This is done by taking multiple “folds” within the datasets as training datasets and taking the mean accuracy over these different validation folds as the cross-validation accuracy^66–68^.

After cross-validation, a Linear Support Vector Machine (SVM) classification method was adopted to classify the behavioral data as the classifier was found to have the best cross-validation output. To further prevent overfitting, a regularization factor was used to impose a penalty on the complexity of the model; this allows simpler classification barriers to be more cost-efficient during training, thus allowing for more generalizable models as long as underfitting has not taken place^69^. For the SVM, a regularization factor of C = 0.1 was utilized during training after hyperparameter tuning between 1.0 and 0.01, while taking order of magnitude steps, showed this value had the highest accuracy while not being overfitted according to the cross-validation metric. Coding was done in both MATLAB (MathWorks, R2022a) and Python (v3.9.7).

## Acknowledgments

We thank Roxanne ter Haar, Mick de Koning and Bram Derksen for assistance with breedings and animal experiments.

## Supplementary Material

**Supplementary figure 1.**
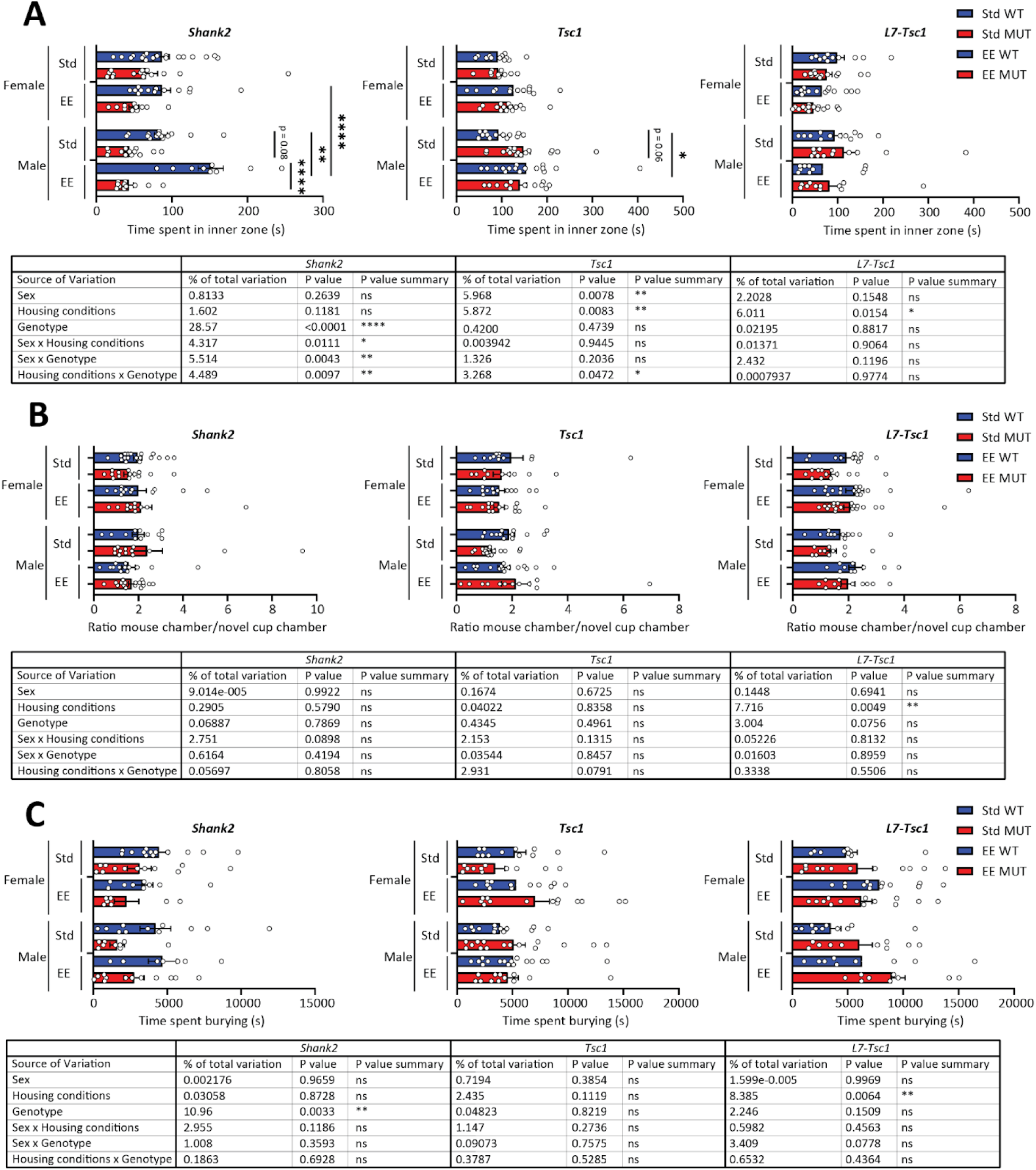
Behavioral phenotyping of autism mouse models using classical behavioral assays. **A)** Time spent in the inner zone of an open field test in Shank2 mice (left), Tsc1 mice (center) and L7-Tsc1 mice (right). Data presented as mean with SEM (three-way ANOVA). **B)** Ratio of the time spent in the chamber containing a novel mouse divided by the time spent in the chamber with a novel object in the social chamber test, in Shank2 mice (left), Tsc1 mice (center) and L7-Tsc1 mice (right). Data presented as mean with SEM (three-way ANOVA). **C)** Time spent burying during the marble burying assay in Shank2 mice (left), Tsc1 mice (center) and L7-Tsc1 mice (right). Data presented as mean with SEM (three-way ANOVA). * p≤0.05, ** p≤0.01, *** p≤0.001

**Supplementary figure 2.**
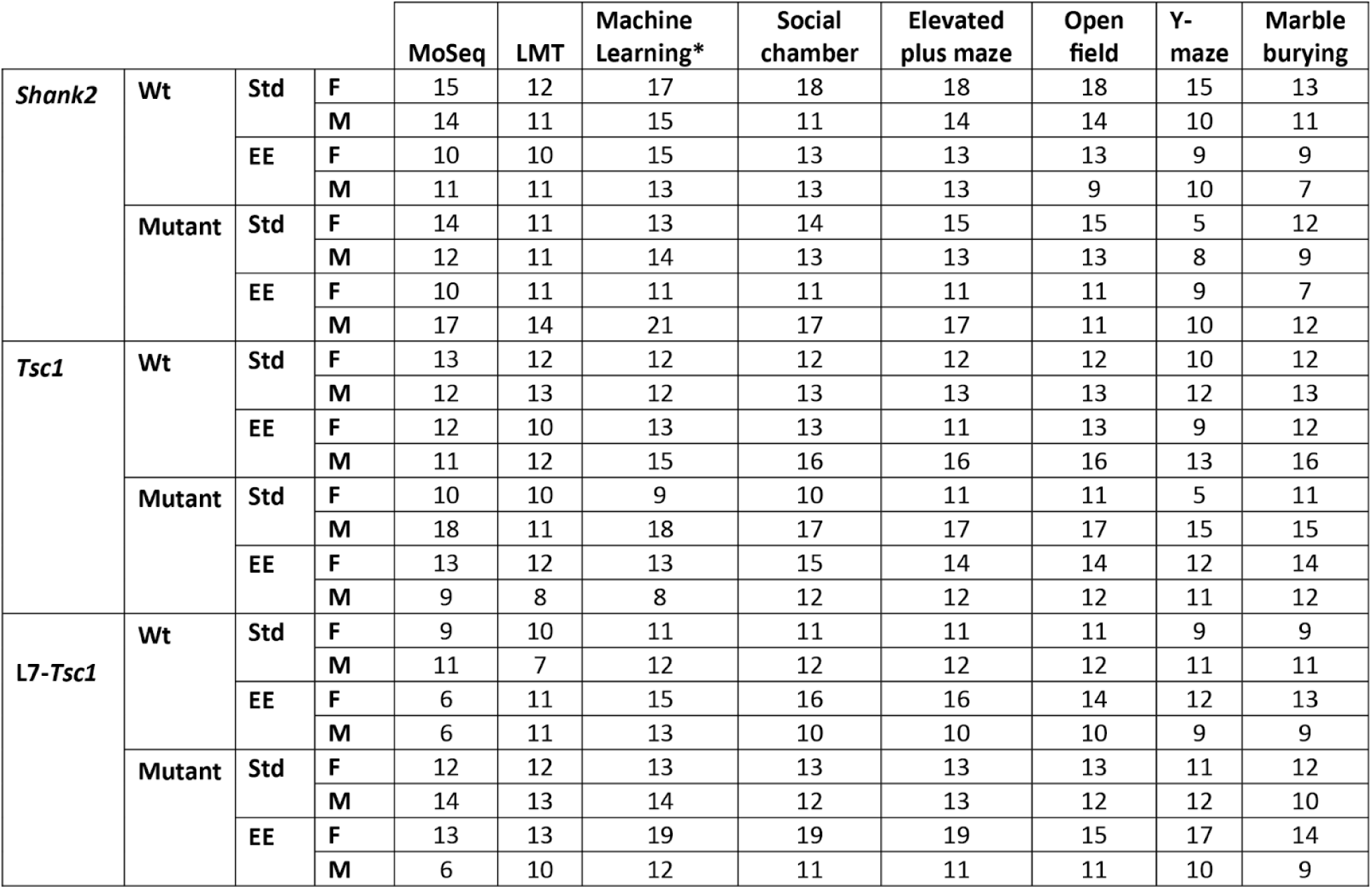
Number of animals used in behavioral testing. The number of animals used per test for classical and multi-parametric behavioral assays. *Data used in dimensionality reduction and machine learning analysis, consisting of data from MoSeq, LMT and single-trait behaviors.

**Supplementary figure 3.**
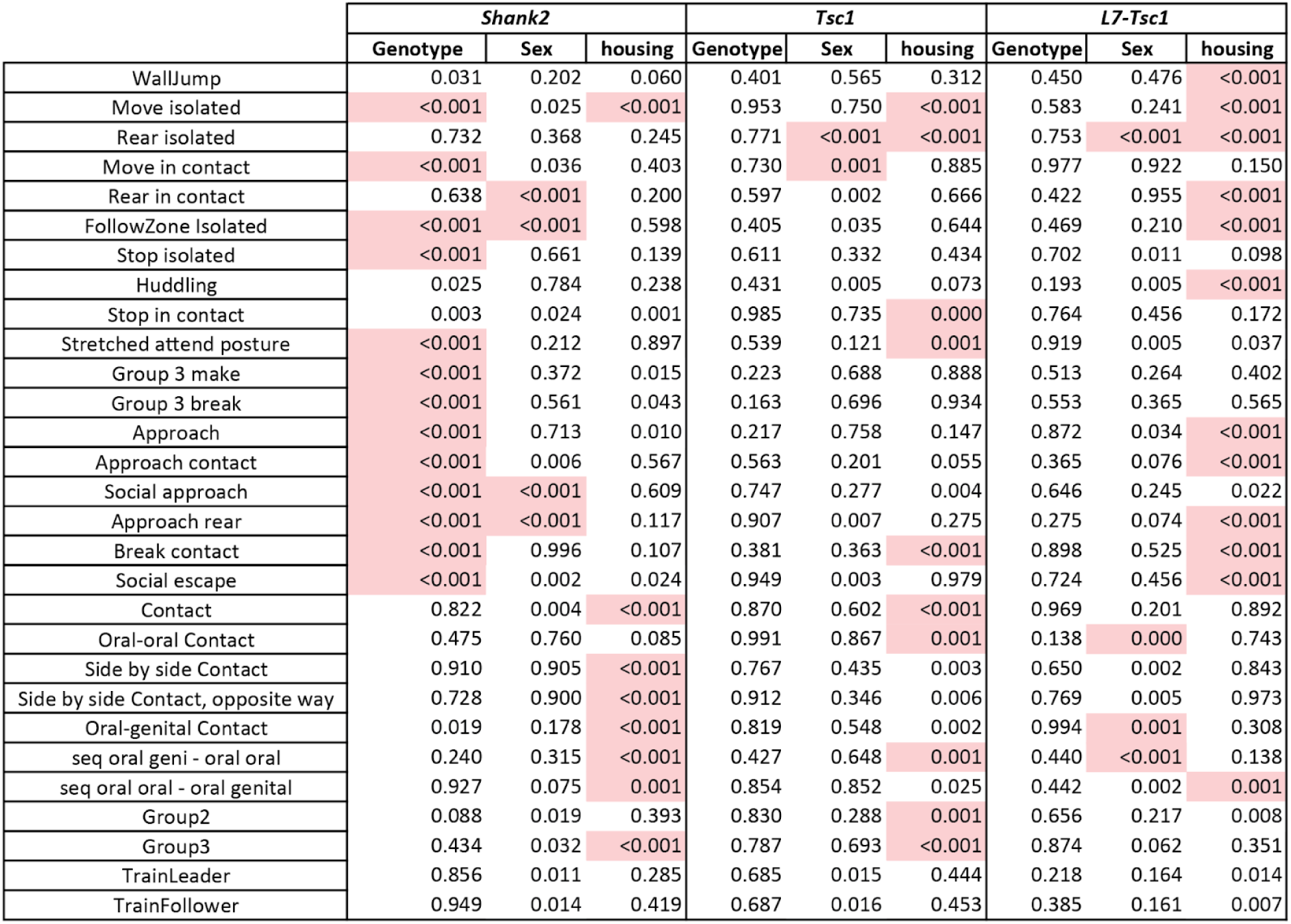
Significance of behaviors in the LMT for sex, housing and genotype differences. Obtained by 3-way anova. A p value of 0.0015 was calculated based on the number of variables and applied to correct for multiple comparisons. Significant cells are highlighted in light red. n = 266 mice.

